# Developing engineering strategies to enhance the genetic stability of fatty alcohol-producing strains for production scale-up

**DOI:** 10.64898/2026.06.01.728301

**Authors:** Jeniffer Perea-Lopez, Yuxin Zhao, Viswanathan Satheesh, Zhanyi Yao, Dan Chen, Zengyi Shao

**Author notes:** corresponding author *Email address:* (Zengyi Shao).

## Abstract

Robust strain stability is essential for industrial-scale fatty alcohol production, as metabolic burden and toxicity not only constrain productivity but also create selective pressure for low-producing or non-producing subpopulations. This genetic instability leads to genetic heterogeneity and compromises strain performance during large-scale fermentation. In this study, we investigated the production stability of fatty alcohol-producing *Yarrowia lipolytica* strains and developed systematic strategies to improve the genetic stability of engineered strains for scale-up production. In a mock fermentation scale-up, a fatty acyl-CoA reductase (*FAR*)-expressing strain lost fatty alcohol production after five consecutive passages. To address this, we fused *FAR* to phosphoglycerate kinase I (*PGK1*), a gene essential to cell growth, to promote the stability of *FAR* expression and prevent production loss. This strategy extended fatty alcohol production by one passage. Additionally, *FAR* was fused to GFP and extended production stability by four additional passages. In parallel, competitive co-culture experiments, in which producing strains were cultured alongside non-producing mutants, revealed that when non-producers emerged with a frequency of 10^*−*5^, they dominated the fermentation population within six passages; at 10%, they took only two passages. Furthermore, deep sequencing of strains that demonstrate different levels of stability and fatty alcohol productivity identified mutation patterns that contribute to strain instability. These findings provide insights into engineering *Y. lipolytica* with enhanced genetic stability for scale-up fatty alcohol production.

## 1. Introduction

Fatty alcohols represent a class of versatile oleochemicals with widespread applications across multiple industries, including cosmetics, pharmaceuticals, lubricants and plastics (Munkajohnpong et al., 2020; Fillet and Adrio, 2016). These amphipathic alcohols serve as building blocks in the synthesis of detergents, emulsifiers and emollients, with global demand continuously expanding (Fortune Business Insights, 2024). Traditionally, fatty alcohols have been derived from petroleum-based feedstocks via the Ziegler process or from bio-based feedstocks (e.g., palm kernel oil) through chemical hydrogenation of fatty acids or fatty acid methyl esters (Su et al., 2024; Shah et al., 2016; Hill et al., 1954; Voeste and Buchold, 1984). These conventional routes, however, have significant environmental drawbacks, including reliance on non-renewable feedstocks and practices that contribute to deforestation, biodiversity loss, and greenhouse gas emissions (Fillet and Adrio, 2016; Fitzherbert et al., 2008). The increasing demand for sustainable and environmentally friendly production methods has driven significant advancements in microbial biosynthesis (Carruthers and Lee, 2022; Scarborough et al., 2022; Adamczyk et al., 2025). Microbial cell factories offer a promising alternative for fatty alcohol production, leveraging metabolic engineering and synthetic biology to enable the development of engineered fatty alcohol-producing microorganisms, including oleaginous yeast, to mitigate the environmental burden of conventional fatty alcohol production methods (Lennen and Pfleger, 2013; Cao et al., 2015; Steen et al., 2010; Zhou et al., 2016; Schultz et al., 2022; Takaku et al., 2020; Xu et al., 2016).

Despite the potential of microbial fatty alcohol production, significant bioprocess constraints impede successful industrial implementation, with strain stability emerging as a critical challenge (Fig. 1). The production of fatty alcohols through heterologous gene expression imposes a substantial metabolic burden on host organisms, diverting cellular resources and energy away from essential metabolic processes (Kastberg et al., 2022; Glick, 1995). Concurrently, the inherent toxicity of fatty alcohols further compromises cellular performance, as fatty alcohol accumulates intracellularly and can intercalate into cellular and subcellular membranes, creating substantial cellular stress and leading to genetic instability and reduced production efficiency during prolonged fermentation processes (Dahlin et al., 2019; Hu et al., 2020; Schultes et al., 2023; Liu et al., 2020). Such stress creates a strong selective pressure that promotes the emergence of genetic variants with reduced metabolic burden, ultimately compromising the genetic stability and production capability of the engineered producing strain (Rugbjerg and Sommer, 2019; Rugbjerg et al., 2018; Mu and Zhang, 2023). These selection dynamics result in genetic heterogeneity within the fermentation population, compromising the consistent performance and economic viability of large-scale bioprocesses (Olsson et al., 2022; Rugbjerg and Olsson, 2020). Consequently, developing robust strategies to maintain genetic stability and mitigate cellular stress represents a critical challenge in engineering fatty alcohol-producing strains.

**Figure 1:**
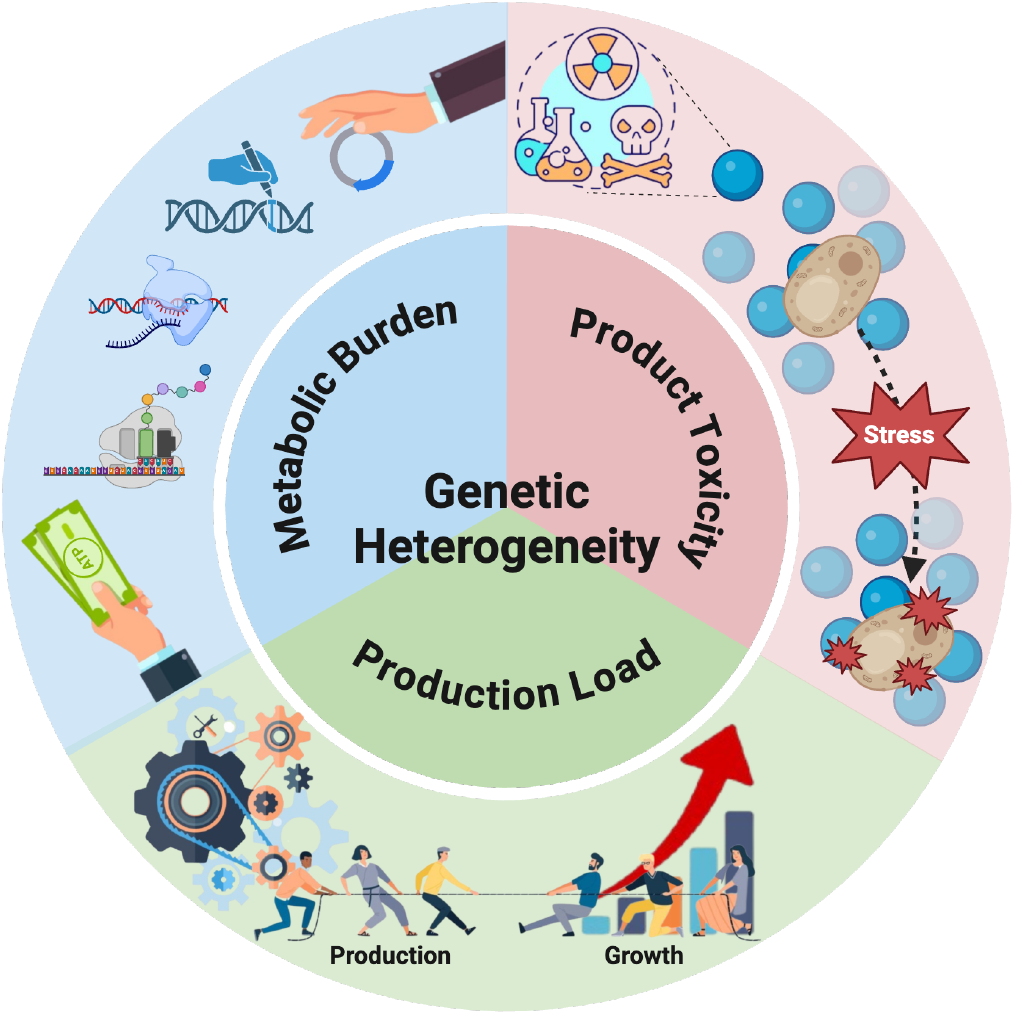
Causal factors underlying genetic heterogeneity in industrial-scale biomanufacturing. There is a bioenergetic cost to introducing additional DNA, whether it be replicating chromosomal or plasmid gene copies, that can result in the depletion of resources (top-left). Additionally, the product could result toxic to the cell and lead to reduced cell fitness (top-right). Furthermore, metabolite-producing pathways may compete with endogenous pathways, imposing a fitness cost by draining cellular energy and central substrates (bottom).

Understanding the genetic mechanisms underlying population heterogeneity in engineered strains can provide critical insights for developing targeted strategies to mitigate genetic instability and optimize long-term strain performance (Rugbjerg et al., 2018, 2021; Munkler et al., 2024). Deep DNA sequencing provides a comprehensive approach for investigating genetic heterogeneity in microbial systems, enabling high-resolution genetic profiling that detects low-frequency variants missed by traditional sequencing methods (Rugbjerg et al., 2021; Wang et al., 2016; Lan et al., 2024). By identifying mutations that arise under selective pressures, this approach offers critical insights into the molecular mechanisms of strain adaptation and genetic instability in metabolic engineering. The technique allows systematic characterization of genetic variations, serving as a tool for subsequent functional analyses to understand their impact on cellular productivity. Prior work has demonstrated the importance of deep sequencing in revealing diverse mutational modes that contribute to genetic heterogeneity, highlighting the potential for early detection of emerging genetic variations (Rugbjerg et al., 2018, 2021). Such approaches facilitate a more targeted and rational approach to developing robust microbial platforms for bioproduction.

In this study, we investigated the genetic stability of fatty alcohol-producing *Yarrowia lipolytica* and developed a novel strain engineering approach to prolong production. We aimed to create a genetic context that would minimize the selective advantage of losing fatty alcohol production capability by directly coupling fatty alcohol production to essential cellular metabolism through the fusion of fatty acyl-CoA reductase (encoded by *FAR*) to phosphoglycerate kinase I (encoded by *PGK1*), and subsequent deletion of the native *PGK1* gene. To comprehensively understand the mechanisms of production-linked genetic instability, we employed a multifaceted approach including serial passaging to simulate production scale-up, deep sequencing of the *FAR* expression cassette to investigate genetic drivers of decreased production, and co-culture experiments to explore the competition between producing and non-producing strains, examining their impact on production stability. By systematically analyzing mutation distributions, population dynamics, and competitive interactions between high-producing and emerging non-producer variants, we sought to develop insights into the evolutionary dynamics of fatty alcohol-producing strains and strategies to mitigate genetic heterogeneity during long-term fermentation.

## 2. Results

### 2.1. Construction of fatty alcohol-producing strains

Fatty alcohol production is both toxic and burdensome on the host strain, compromising cell viability and reducing overall fatty alcohol productivity, thereby posing a significant challenge to their feasibility for industrial-scale bioproduction (Su et al., 2024). We previously engineered the fatty alcohol production pathway in *Yarrowia lipolytica* by integrating a single copy of *Marinobacter aquaeolei* VT8 *FAR* into the *AXP* locus, a known neutral integration site. To assess the production stability of the Δ*AXP::FAR* strain, we conducted a mock test simulating fermentation scale-up, continuously passaging cells from the previous fermentation culture into fresh growth medium and using those cells to start a new fermentation culture. We quantified the amount of fatty alcohol produced, primarily palmityl alcohol (C16:0), stearyl alcohol (C18:0), and oleyl alcohol (C18:1), for each passaged culture and observed the loss of fatty alcohol productivity after five consecutive passages. Cell density was also measured at the end of the fermentation period to assess potential variations in growth and determine whether any differences in fatty alcohol production were due to changes in cell growth rates across passages. We observed no significant changes in cell density across passages, suggesting that the decline in fatty alcohol production was not due to reduced cell growth. Instead, this decline likely reflects genetic instability within the production pathway, leading to the accumulation of mutations that disrupt *FAR* expression and consequently reduce fatty alcohol biosynthesis. To address the issue of production-linked genetic instability and impose a selective pressure that favors fatty alcohol biosynthesis, we aimed to develop a growth-coupled fatty alcohol production pathway, ensuring that sustained production provides a competitive advantage to the host strain.

To circumvent production-linked genetic instability, we fused *FAR* to a monomeric essential gene, thereby creating a metabolic dependency whereby cells must maintain *FAR*, and thus fatty alcohol production, to remain viable. We hypothesized that this protein-fusion strategy would impose a selective pressure against frameshift mutations and other disruptions in *FAR*, as such mutations would concurrently compromise the function of the essential gene. As a result, cells would be driven to retain an intact *FAR*, promoting the long-term stability of fatty alcohol production. Glycolysis presents an ideal target for growth-production coupling, as it is responsible for energy generation and precursor supply for numerous cellular processes, including fatty acid biosynthesis (Kierans and Taylor, 2024). Previous work has demonstrated this concept of coordinating fatty alcohol biosynthesis with glycolysis through *FAR* expression under glycolytic promoters, creating a ‘push-and-pull’ effect that enhanced production (Zhang et al., 2019). Thus, our approach of fusing *FAR* to a glycolytic gene may create a beneficial ‘push-and-pull’ effect, where glycolytic flux drives fatty acid precursor availability while FAR simultaneously directs intermediates towards fatty alcohol biosynthesis, potentially alleviating feedback inhibition and enhancing production efficiency. Among the glycolytic enzymes, PGK1 emerged as a suitable candidate for a protein-fusion partner due to its monomeric nature, lack of oligomerization and moderate size, minimizing structural disruptions and reducing the likelihood of steric hinderances that could affect the fused-protein function and stability.

In this study, we engineered two distinct fatty alcohol-producing strains in *Y. lipolytica* using the Lowered Indel Nuclease system Enabling Accurate Repair (LINEAR) CRISPR platform for homologous recombination (HR)-based genome editing (Ploessl et al., 2022) (Fig. 2A, Supplementary Table S1). The first strain implemented our proposed growth-production coupling strategy, wherein *FAR* was fused to *PGK1* via a flexible (GGGGS)_3_ protein linker. We initially attempted a single-step approach to simultaneously integrate the *FAR-PGK1* fusion construct while knocking out the endogenous *PGK1* gene by targeting the *PGK1* locus directly. Despite modifying the N20 sequence within the fused *PGK1* gene to avoid guide RNA recognition, we were unable to achieve successful *PGK1* knock-out in a single step. Consequently, we adopted a two-step process: first integrating the *FAR-PGK1* fusion construct from the previous approach into the neutral *AXP* integration site, followed by targeted deletion of the endogenous *PGK1* gene, wherein the *URA3* selection marker was integrated, thereby linking fatty alcohol production to an obligate metabolic pathway and yielding the Δ*AXP::FPG* strain. The second strain, Δ*AXP::FGU*, served as a control, featuring *FAR* fused to GFP via the (GGGGS)_3_ protein linker and integrated into the *AXP* locus, enabling visual monitoring of *FAR* expression and stability throughout cultivation.

**Figure 2:**
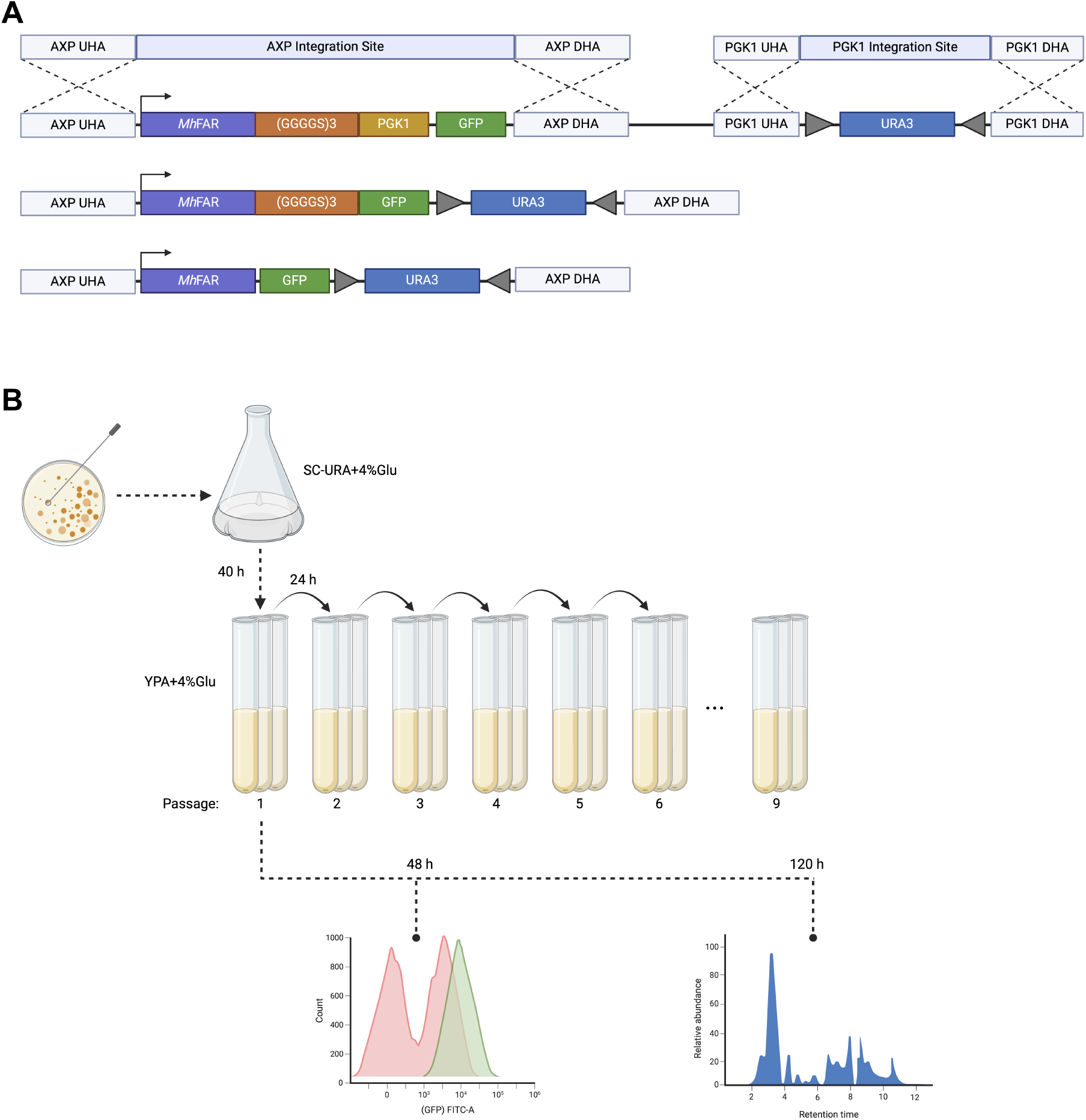
Construct design and experimental workflow for assessing fatty alcohol production stability. A) Schematic representation of the three engineered constructs in *Y. lipolytica*. B) Scale-up cultivation of strains was experimentally simulated by serial passaging of growing cultures and subjected to flow cytometric analysis to study population dynamics and GC-MS analysis to assess production stability.

### 2.2. Assessment of fatty alcohol production stability in engineered strains

Recent studies in yeast and bacteria have employed serial passaging strategies to simulate industrial scale-up and long-term cultivation conditions (Rugbjerg et al., 2018, 2021; Lv et al., 2020; D’Ambrosio et al., 2020). This experimental design effectively simulates industrial stability assessments, providing a reasonable approximation of the challenges encountered during commercial scale-up processes. Following confirmation that the cellular growth of the Δ*AXP::FPG* strain had not been negatively impacted, we proceeded to assess the stability of the fatty alcohol production pathway in the engineered strain, with the Δ*AXP::FAR* and Δ*AXP::FGU* strains as controls. To simulate fermentation scale-up, we serially passaged fatty alcohol-producing cells strictly in the exponential growth phase every 24 h to avoid selecting for stationary-phase culture transitions and performed nine sequential passages (Fig. 2B). Here, seed cultures were initially grown in SC-URA+4% Glu medium to select for the URA3 auxotroph and then transferred into YPA+4% Glu medium for fermentation. For the first passage, the seed cultures were inoculated into nonselective medium at an OD_600_ of approximately 0.2. For each of the following passages, cultures were subjected to a 1:100 dilution into fresh medium every 24 h followed by a 120 h fermentation.

To monitor the fermentation population and detect the emergence of potential non-producing subpopulations, samples were collected for flow cytometric analysis at 48 h after each passage (Fig. 3A). The Δ*AXP::FGU* strain enabled direct monitoring of production strain integrity, as GFP fluorescence is contingent upon correct *FAR* expression. Any frameshift mutations or other mutations disrupting *FAR* could consequently abolish GFP fluorescence, allowing real-time detection of emerging non-producing subpopulations. For both the Δ*AXP::FAR* and Δ*AXP::FPG* strains, GFP fluorescence remained relatively stable, as GFP is independent of *FAR* expression. In contrast, for the Δ*AXP::FGU* strain, we observed a decline in the GFP-expressing population by the sixth passage along with the emergence of a non-expressing subpopulation after the fourth passage, steadily increasing and eventually overtaking the fermentation population. This finding suggests that GFP expression is dependent on *FAR* expression, enabling the tracking of genetic instability within each passage. The observed decline in GFP fluorescence and the emergence of a non-expressing subpopulation indicate that mutations disrupting *FAR* activity may be preferentially selected over time, as such mutations lead to the loss of GFP fluorescence. Consequently, a corresponding decline in fatty alcohol production would also be expected, given the direct linkage between *FAR* expression and GFP fluorescence. Having assessed the genetic instability of the engineered strains across various passages via flow cytometry, we next sought to examine the stability of the fatty alcohol production pathway in relation to productivity by measuring total fatty alcohol titer at the conclusion of each passage (Fig. 3B). Samples were collected at the end of each 120 h fermentation for each passage, and the fatty alcohol titer was quantified by measuring the amounts of C16:0, C18:0 and C18:1 fatty alcohol produced. Results showed distinct profiles of fatty alcohol production across the different constructs. For the Δ*AXP::FAR* strain, fatty alcohol production peaked on the fourth passage, reaching a titer of 346 mg/L, before falling to negligible levels by the fifth passage. TheΔ*AXP::FPG* strain similarly achieved its highest output at the fourth passage but attained a substantially higher titer of 612 mg/L. Subsequently, production was halved by the fifth passage and declined to minimal levels by the sixth passage. In contrast, the Δ*AXP::FGU* strain exhibited a different pattern, with production steadily increasing through the first five passages and reaching a maximum titer of 440 mg/L at the fifth passage. This was followed by a gradual decrease in fatty alcohol production until the seventh passage, after which a rapid decline occurred, resulting in negligible titers by the ninth passage. This production profile correlates with our flow cytometric analysis, which revealed a decline in the GFP-expressing population at the sixth passage, with a non-expressing subpopulation progressively dominating the fermentation population. As the onset of this population shift coincided with the observed decrease in fatty alcohol production, the emergence of this non-expressing subpopulation may represent a non-producing variant in which disruption in *FAR* expression simultaneously led to loss of GFP fluorescence, explaining both the declining fatty alcohol production and the shift in population fluorescence observed in later passages. Overall, these results indicate that the protein-fusion strategy between the essential PGK1 and FAR extended the stability of fatty alcohol production by one additional passage compared to the FAR-only strain. More notably, fusing FAR to GFP significantly enhanced production stability, extending production capability by four additional passages.

**Figure 3:**
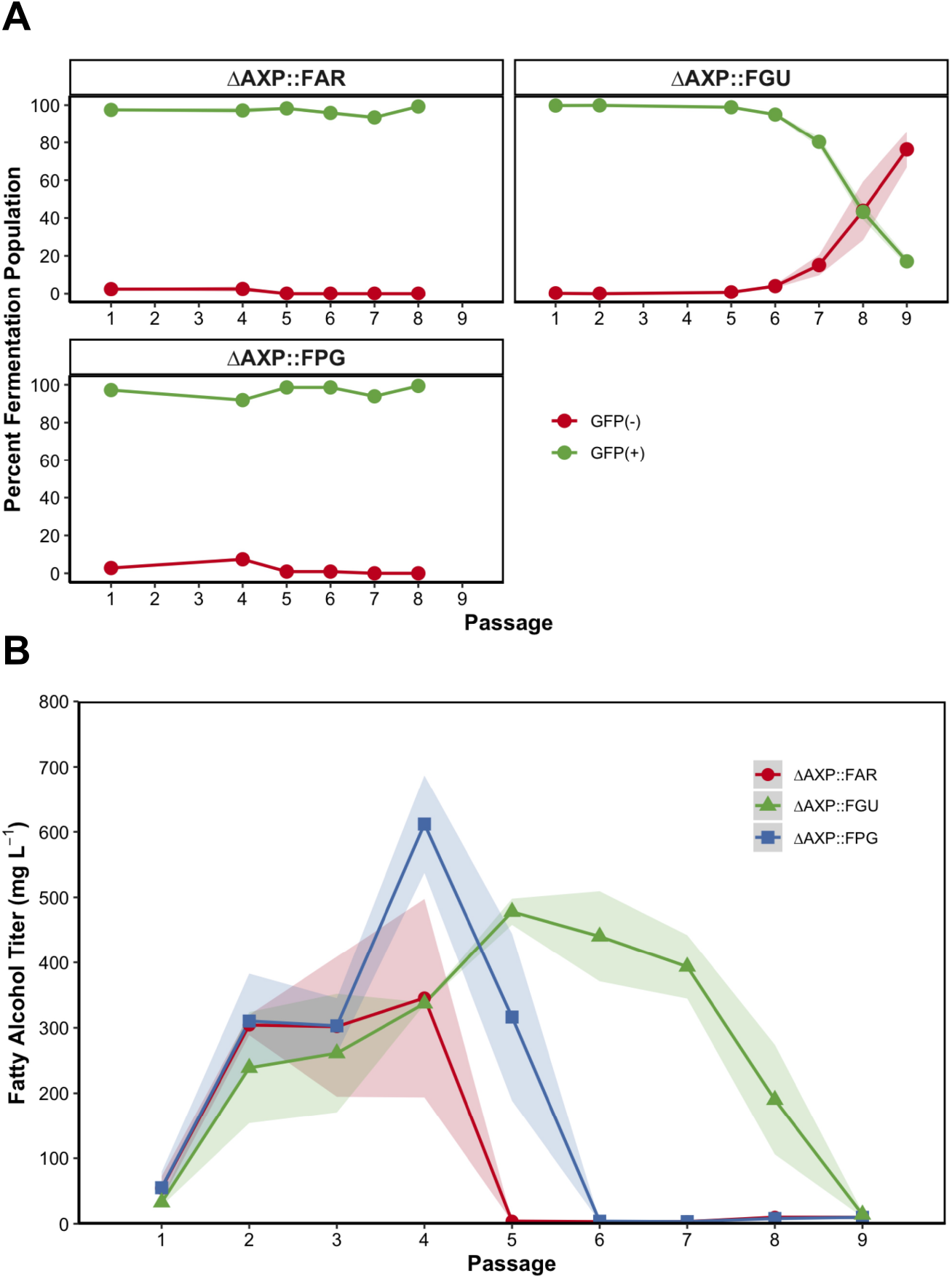
Population dynamics and production stability of engineered fatty alcohol-producing *Y. lipolytica* strains. A) Flow cytometric analysis of engineered strains (green) after 48 h of each passage to monitor fermentation population and detect emerging non-producing strains (red). B) GC-MS analysis of the fatty alcohol titers for the passaged cultures after 120 h fermentation. For all sample points, the mean is shown with error bars representing standard deviation of the mean from three (n=3) biological replicates.

### 2.3. Deep sequencing to elucidate production-linked genetic instability

To investigate whether the observed phenotypic decline in fatty alcohol production stemmed from genetic heterogeneity and elucidate the underlying causes of genetic instability, we conducted deep sequencing of the integration site of the three engineered fatty alcohol-producing *Y. lipolytica* strains, which had exhibited varying levels of stability and fatty alcohol productivity over the course of successive passages (Fig. 4). Genomic DNA was isolated from cultures at each passage, beginning at the first passage and continuing until fatty alcohol production became negligible. For each strain and passage, we pooled the three parallel-cultivated populations in equal proportions before DNA extraction. We then amplified the integration site containing the *FAR* expression cassettes to analyze the distribution of mutations within this region. The amplicons were sequenced using short pair-end reads (2×250 base pair Illumina sequencing) with overall sequencing coverage generally ranging between 10,000 to 30,000 reads per base and a quality threshold of Q30.

**Figure 4:**
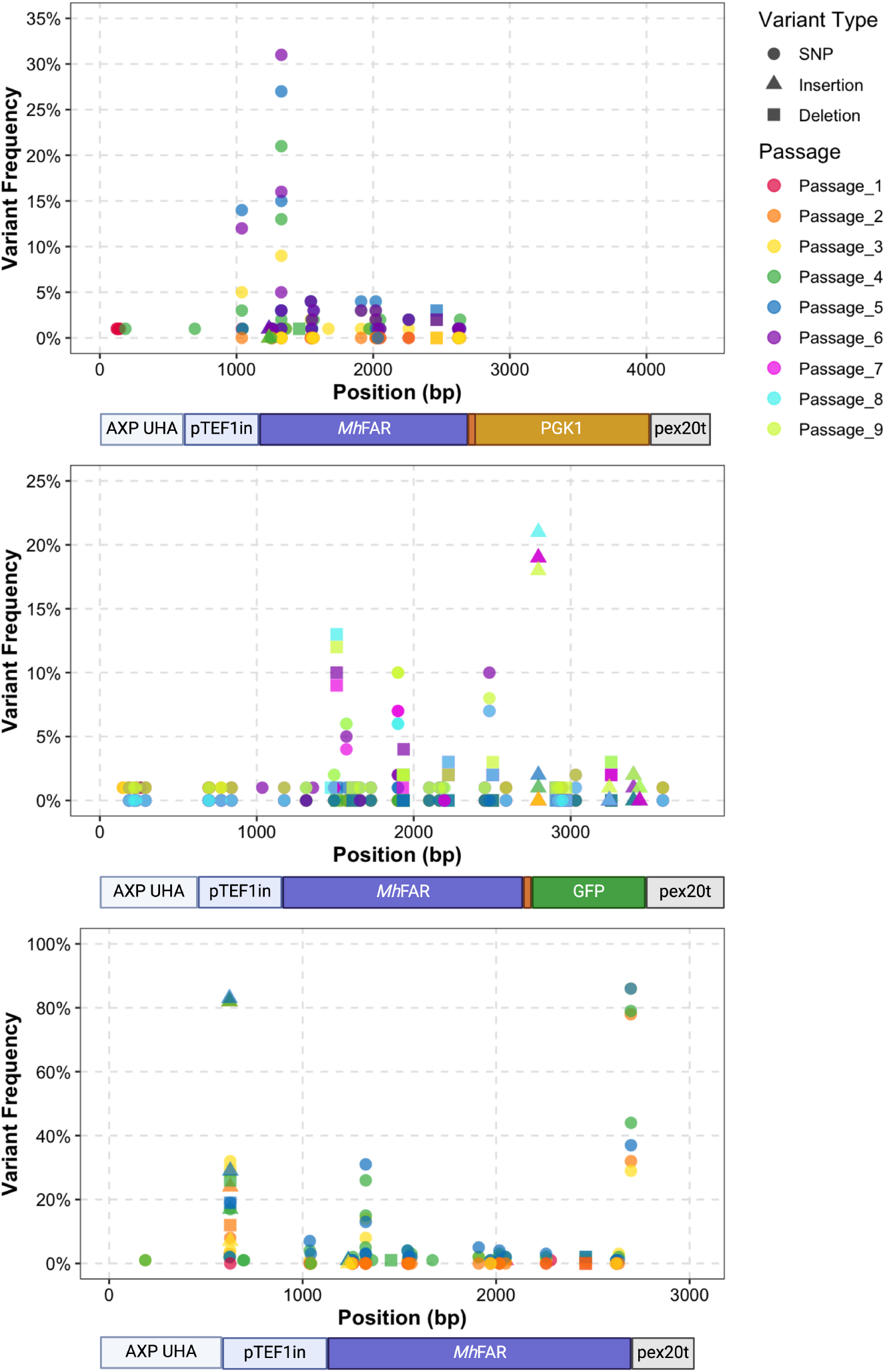
Mapping mutation patterns via deep sequencing to investigate decreased fatty alcohol production. Variants were filtered using a 1% frequency threshold with consideration of low-frequency variants from earlier passages. Shapes indicate variant type and colors represent individual passages. The underlying schematic illustrates the genomic regions sequenced, providing spatial context for observed genetic variations.

To analyze genetic variations, we calculated the variant frequencies by dividing the variant count by the total count at each position. We established a 1% frequency threshold for variant inclusion, while also incorporating variants below this threshold from earlier passages if they corresponded to filtered variants in the final passage. We examined variants within the *FAR* expression cassette and the upstream region and identified several variants that exhibited significant frequency in the final passage. In the Δ*AXP::FAR* strain, we identified multiple single-nucleotide polymorphisms (SNPs), including a SNP at codon 57 of the *FAR* gene, resulting in an arginine substitution with leucine (G1326T; 31%), glutamine (G1326A; 13%) or proline (G1326C; 3%). Additionally, a SNP in the stop codon resulted in a mutation, changing the stop codon to glycine (T2696G; 86%). These SNP variants increased in frequency across several passages concurrent with declining fatty alcohol production. For the Δ*AXP::FPG* strain, we did not observe any frameshift mutations in *FAR* nor variants in *PGK1*. We did, however, observe two SNPs–one in the promoter region (G1039A; 12%) and another at codon 57 of the *FAR* gene previously observed in the Δ*AXP::FAR* strain which resulted in an arginine-to-leucine substitution (G1328T; 31%). The Δ*AXP::FGU* strain exhibited a single-base deletion in codon 118 of the *FAR* gene (1509delG; 12%), causing a frameshift and altered downstream amino acid sequence and resulting in premature termination. We also detected a SNP in the FAR active site that resulted in a tyrosine-to-phenylalanine substitution at residue 248 (A1899T; 10%), a change known to inactivate FAR (Filling et al., 2002). Other prevalent variants identified were a SNP in codon 442 of the *FAR* gene (T2482G; 8%), resulting in a tyrosine-to-stop codon mutation, and a single-base insertion between codons 18 and 19 of GFP (2794 2795insA; 8%), causing a frameshift mutation that resulted in the stop codon being positioned later in the sequence, leading to an extended protein. When comparing the results from deep sequencing with fatty alcohol production in the Δ*AXP::FGU* strain, we noted a sudden increase in variants in the sixth and seventh passage, particularly those aforementioned. This genetic shift coincided with the decline in fatty alcohol production observed after the fifth passage, providing a link between genetic instability and the observed production loss phenotype.

### 2.4. Colony-level analysis to evaluate production-loss variants

To determine whether the variants revealed by deep sequencing were co-occurring, we performed a colony-level genetic analysis. For this study, we focused on the Δ*AXP::FGU* strain, as it exhibited the most stability in fatty alcohol production. From the final passage (i.e., Passage 9), we isolated 60 individual colonies and amplified the integration site. Sanger sequencing of the fusion region enabled us to detect the variants identified in our deep sequencing results and assess their potential co-occurrence within individual colonies. We specifically examined variants that had been observed at high frequencies, reasoning that their prevalence suggested they should also be detectable within individual colonies. Co-occurring variants were largely absent, with the exception of two SNPs within the FAR active site, which together would result in a loss-of-function mutation. However, when these variants occur separately, only the SNP at position 1899 is sufficient to inactivate FAR, as the T1900C mutation is a synonymous substitution that does not alter the encoded amino acid. While our analysis prioritized high-frequency variants due to their greater likelihood of detection within individual colonies, it remains possible that additional co-occurring variants were present at lower frequencies and not captured in our sample set.

To assess whether the observed variants conferred a growth advantage, we performed a growth assessment on 30 individual colonies over 120 h, comparing them to the wild-type Δ*AXP::FGU* strain (Fig. 5A). Given the reduction in fatty alcohol production observed in the ninth passage, we sought to determine whether the reduced production would result in improved growth. During the exponential phase, the variants exhibited a higher growth rate than the wild-type strain, suggesting that the loss or reduction of fatty alcohol production alleviated the metabolic burden or toxicity on the host cell. At the end of the 120 h fermentation, we quantified fatty alcohol production and found that most variants had lost the ability to produce fatty alcohol (Fig. 5B). These findings, coupled with the observed growth advantage, indicate that the disruption of fatty alcohol biosynthesis likely contributed to the improved growth observed. Notably, a few variants retained fatty alcohol production and exhibited slightly higher titers than the wild-type strain (strains P9FG42 and P9FG54), suggesting that these variants may have evolved to achieve both enhanced growth and sustained production. Additionally, we did not observe any high-frequency variants from our colony-level genetic analysis in these variants, particularly those within the *FAR* gene, that would result in a loss-of-function mutation.

**Figure 5:**
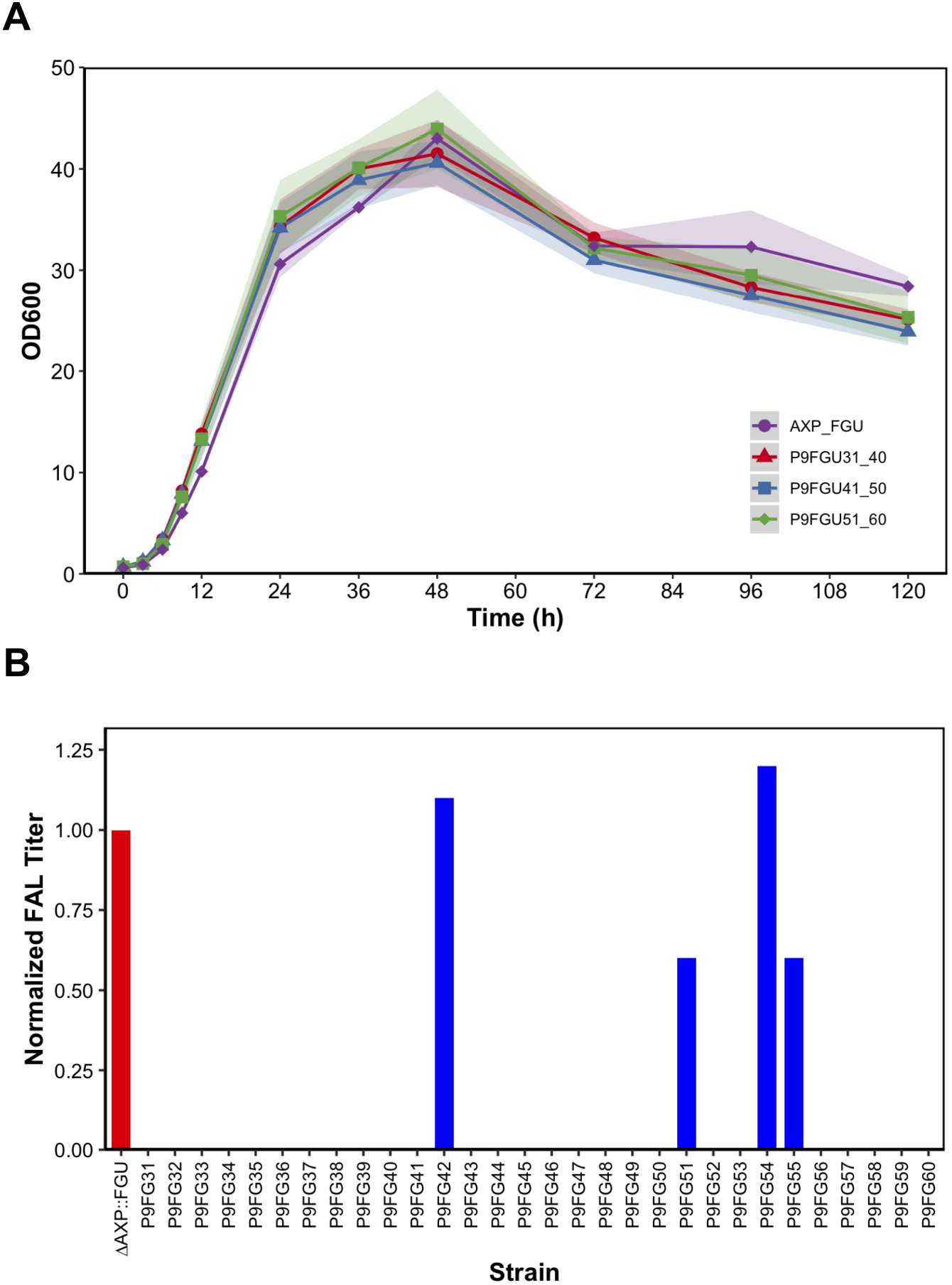
Assessment of Δ*AXP::FGU* variants from Passage 9 . A) Growth assessment of colonies 31 through 60 (strains P9FGU31-60). B) Fatty alcohol titers were measured and normalized to the wild-type strain (red) to assess production across the colony variants.

Next, we aimed to determine whether the strains P9FG42 and P9FG54, which exhibited improved fatty alcohol production and a higher growth rate compared to the wild-type strain, had evolved to be more stable. Given their enhanced performance, we hypothesized that these strains may have undergone genetic changes contributing to both increased growth and production stability. To assess the genetic stability, we serially passaged these strains following the previously described protocol. Flow cytometric analysis was performed at 48 h, and fatty alcohol production was quantified after a 72 h fermentation, using strains Δ*AXP::FAR* and Δ*AXP::FGU* as controls. Flow cytometric analysis revealed a decline in the GFP-expressing population in strain P9FG42 after the fourth passage, accompanied by the emergence of a non-expressing subpopulation (Fig. 6A). In contrast, strains Δ*AXP::FGU* and P9FG54 exhibited similar population dynamics, with this shift occurring after the sixth passage. At the conclusion of the 72 h fermentation, we assessed the production stability for each consecutive passage and observed that fatty alcohol production steadily declined in strains P9FG42 and P9FG54, with significantly reduced titers observed by the ninth passage—comparable to the wild-type strain (Fig. 6B). Despite retaining production up until the ninth passage, the decline in both fatty alcohol production and GFP-expressing populations in strains P9FG42 and P9FG54 mirrors the patterns observed in the wild-type strain, indicating that these strains did not evolve increased stability. Furthermore, this suggests that, like the wild-type strain, these strains may be susceptible to the same genetic or regulatory factors that ultimately reduce production, even in strains that initially exhibited improved performance.

**Figure 6:**
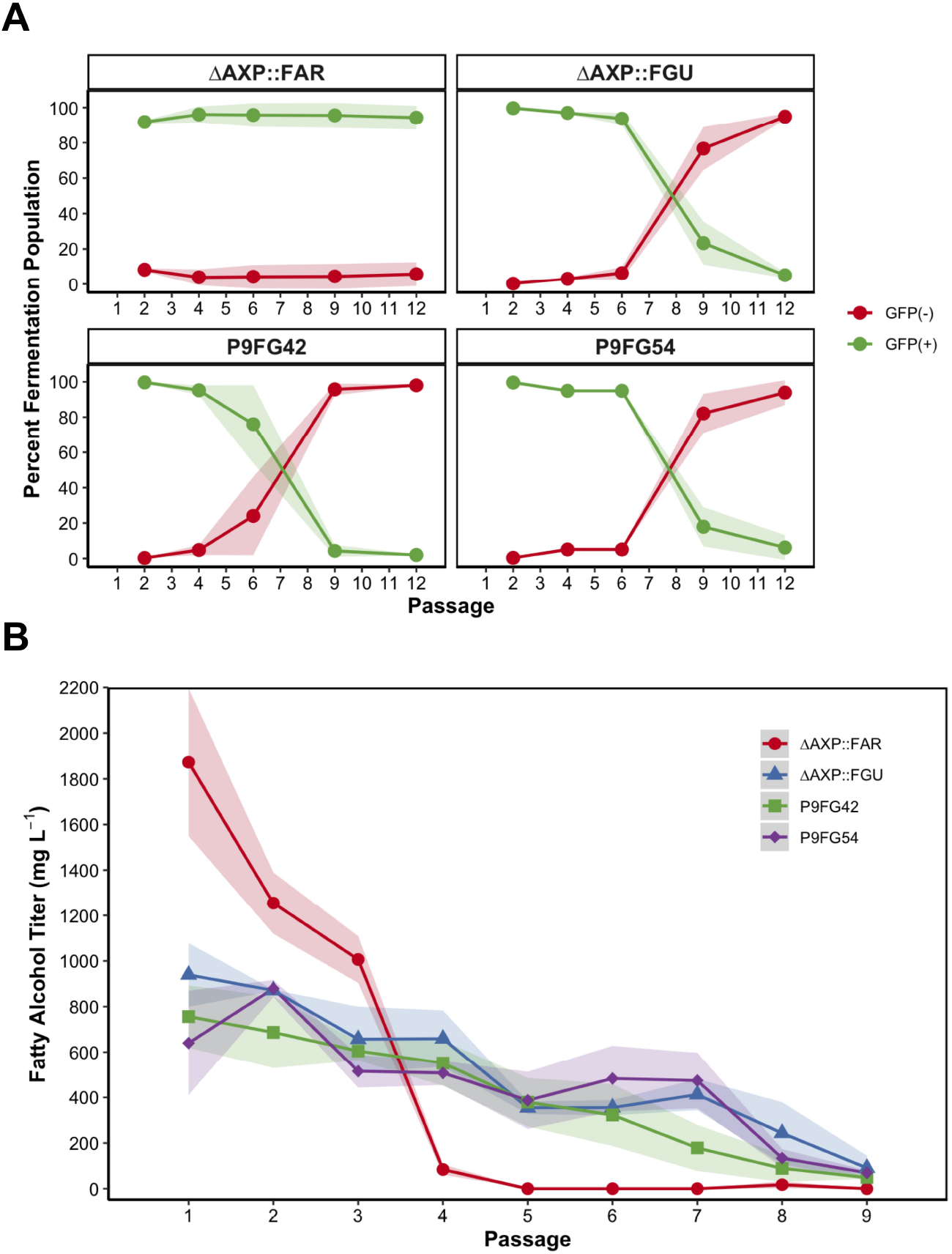
Population dynamics and production stability of fatty alcohol-producing variants P9FG42 and P9FG54. A) Flow cytometric analysis of the variant strains, including Δ*AXP::FAR* and Δ*AXP::FGU* as controls, monitored during fermentation to assess the emergence of a non-producing population. Data show the distinct distribution of producing (green) and non-producing (red) cells in the population. B) Fatty alcohol production for the consecutive passages over a 72 h fermentation period to evaluate the stability of fatty alcohol production. For all sample points, the mean is shown with error bars representing standard deviation of the mean from three (n=3) biological replicates.

### 2.5. Analyzing genetic and extragenic factors of production heterogeneity

While deep sequencing provided valuable insights into genetic variants and their emergence patterns across successive passages, the specific impacts of these variants on protein structure and fatty alcohol productivity warranted further investigation. To functionally characterize these mutations, we designed synthetic mutants based on the high-frequency variants identified through deep sequencing of the three engineered strains. Each construct was modeled after the FAR-GFP fusion construct, with individual mutations introduced into the promoter, *FAR*, or GFP region. Rather than creating constructs with co-occurring variants, we systematically tested each variant separately to isolate its specific effects. For the mutated stop codon observed in the Δ*AXP::FAR* strain, this change was incorporated into the FAR-only construct. All engineered constructs were then integrated into the AXP locus of *Y. lipolytica* to directly compare how each individual variant influenced fatty alcohol productivity relative to the wild-type Δ*AXP::FGU* strain.

Following a 72 h fermentation, we quantified fatty alcohol production in the synthetic mutant strains and compared their productivity to the wild-type Δ*AXP::FGU* strain, which produced 133.4 mg/L (Fig. 7). Among the six mutants strains, three exhibited substantially reduced production: FAR(G1039A)-GFP (promoter SNP), FAR(G1326C)-GFP (arginine-to-proline substitution) and FAR-GFP(2794 2795insA) (frameshift mutation resulting in delayed stop codon). On average, these strains produced titers less than 30 mg/L. Strains FAR(G1326T)-GFP and FAR(G1326A)-GFP, which introduced arginine-to-leucine and arginine-to-glutamine substitutions, yielded 76.2 mg/L and 112.0 mg/L, respectively. Notably, strain FAR(T2696G), which introduced a stop codon-to-glycine mutation, produced 281.7 mg/L—more than twice the titer of the wild-type strain. These findings highlight the critical role of specific genetic variants in modulating fatty alcohol biosynthesis. While most mutations led to reduced production, the stop codon-to-glycine mutation unexpectedly resulted in more than a twofold increase in fatty alcohol titer, suggesting that extended translation may enhance enzyme activity. Notably, in the Δ*AXP::FAR* strain, the emergence of this mutation initially coincided with increased fatty alcohol production but was later associated with a rapid decline in productivity. Further stability assessments are needed to determine whether this enhancement resulted in the genetic instability and eventual loss of productivity observed.

**Figure 7:**
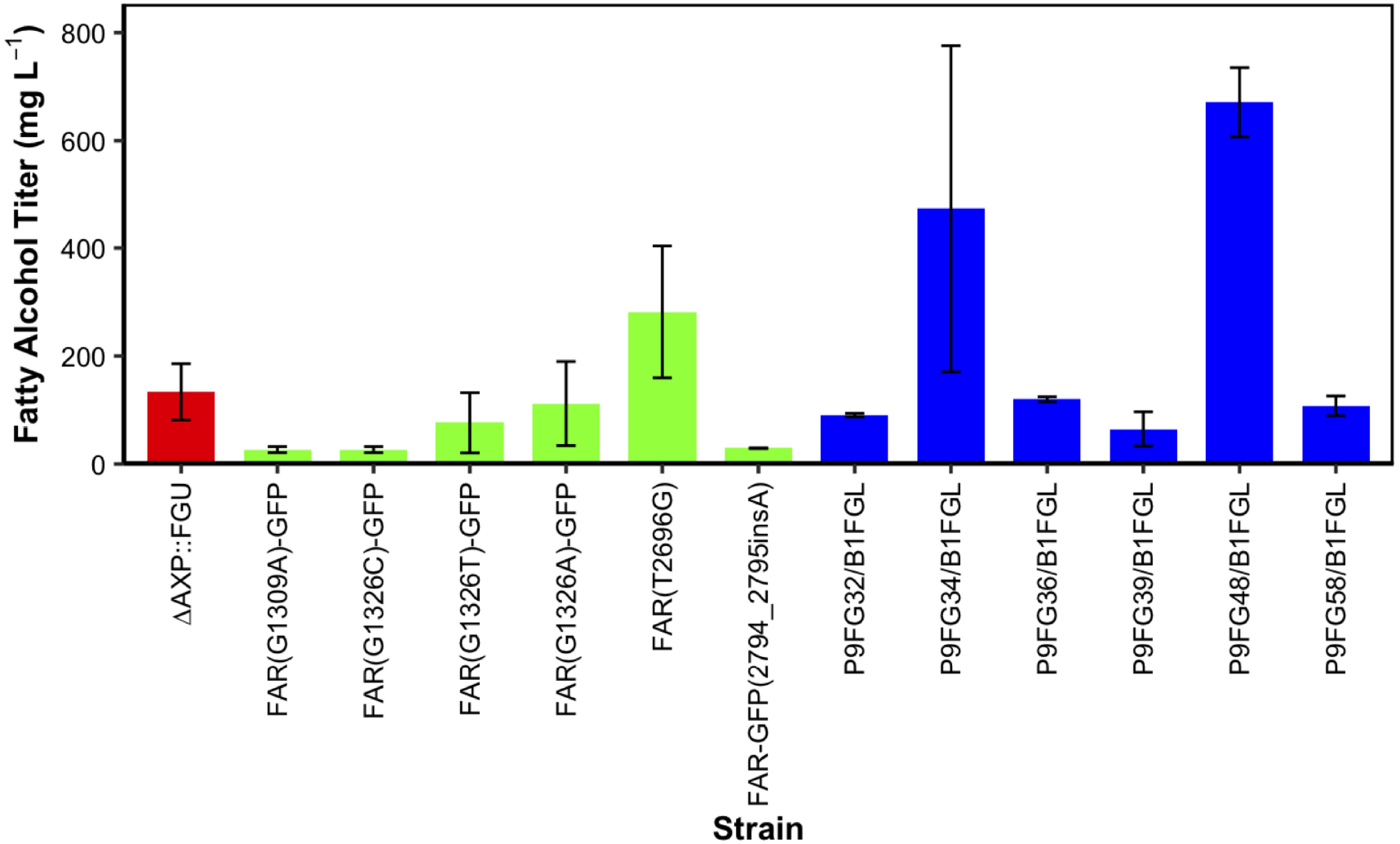
Genetic variability and restoration of fatty alcohol production in engineered and mutant strains. Fatty alcohol production in synthetic mutants (green) and strains with restored production (blue), compared to the wild-typeΔ*AXP::FGU* strain (red) after a 72 hours fermentation. For all sample points, the mean is shown with error bars representing standard deviation of the mean.

While our findings focused on the variants observed within the *FAR* expression cassette, we did not explore potential mutations outside of this region or other mechanisms that may have inhibited fatty alcohol production. To address this, we aimed to restore fatty alcohol production in strains that had lost the ability to produce. We hypothesized that mechanisms inhibiting fatty alcohol production would still be present even when a functional *FAR* cassette is integrated into another locus. Thus, we integrated a functional *FAR-GFP* cassette into the *B1* locus, a stable integration site. From our previous colony-level production analysis, we selected six strains that exhibited production loss. From the colony-level genetic analysis of these strains, we found that three strains (P9FG32, P9FG34, P9FG36) had no detectable mutations, with no SNPs or indels identified. In contrast, we observed mutations in the other three strains: P9FG39 had a A1899T mutation in the *FAR* gene, resulting in an inactive enzyme; P9FG48 contained a 1510delG mutation, leading to a frameshift and an early stop codon; and P9FG58 had a 2794 2795insA mutation in GFP, causing a frameshift and delayed stop codon.

After a 72 h fermentation, all strains successfully produced fatty alcohols at levels comparable to the wild-type Δ*AXP::FGU* strain (Fig. 7). Notably, strains P9FG34/B1FGL and P9FG48/B1FGL exhibited significantly enhanced fatty alcohol production, yielding 473.2 mg/L and 671.2 mg/L, respectively—more than fourfold the wild-type titer. These results suggest that while some mutations may have directly disrupted FAR function, production loss in certain strains was likely due to regulatory or extragenic factors rather than irreversible genetic disruptions within the *FAR* cassette itself. The ability to restore and enhance production at an alternative locus highlights the influence of genomic context on production stability, warranting further investigation into mechanisms beyond the *FAR* expression cassette and mutagenesis.

### 2.6. Competitive dynamics and the role of genetic heterogeneity in fatty alcohol production

Building on our previous analysis of genetic heterogeneity and its impact on fatty alcohol production, we sought to further explore the competitive dynamics between producing and non-producting strains within a fermentation culture. To simulate the population dynamics of the fermentation culture and investigate the competition between fatty alcohol-producing strains and emerging non-producing variants, we constructed a synthetic mutant in which an inactivated *FAR* was fused to a codon optimized blue fluorescent protein (*Yl* mTagBFP2) (Subach et al., 2011). To generate a loss-of-function mutation in *FAR*, we substituted tyrosine 248 with phenylalanine (Y248F), disrupting its catalytic activity. A 120 h growth assessment revealed no significant differences in growth between the producing Δ*AXP::FGU* strain and the non-producing Δ*AXP::F*BU* strain. Using the same serial passaging protocol as before, we co-cultured the producer strain and the non-producer strain at various initial seeding ratios in the first passage (Fig. 8A). The producer strain was inoculated at an OD_600_ of approximately 0.2 with producer-to-nonproducer (GFP-to-BFP) ratios of 1:0 (producer only), 10:1, 100:1, 1000:1 and 10,000:1. Flow cytometric analysis and fatty alcohol quantification were performed across successive passages to assess strain dynamics and production stability. Flow cytometric analysis revealed that the non-producer strain became dominant in the fermentation culture by the second passage in the 10:1 condition and by the sixth passage in the 10,000:1 condition (Fig. 8B). Fatty alcohol quantification further demonstrated that higher initial fractions of the mutant subpopulation correlated with a more rapid loss of production (Fig. 8C). Specifically, cultures seeded at a 10:1 GFP-to-BFP ratio lost production after four passages, whereas the 10,000:1 condition maintained stable production until approximately the ninth passage, similar to the wild-type Δ*AXP::FGU* strain. These findings underscore the profound impact of genetic heterogeneity on fatty alcohol productivity, revealing how the emergence of non-producing variants can rapidly compromise production capability.

**Figure 8:**
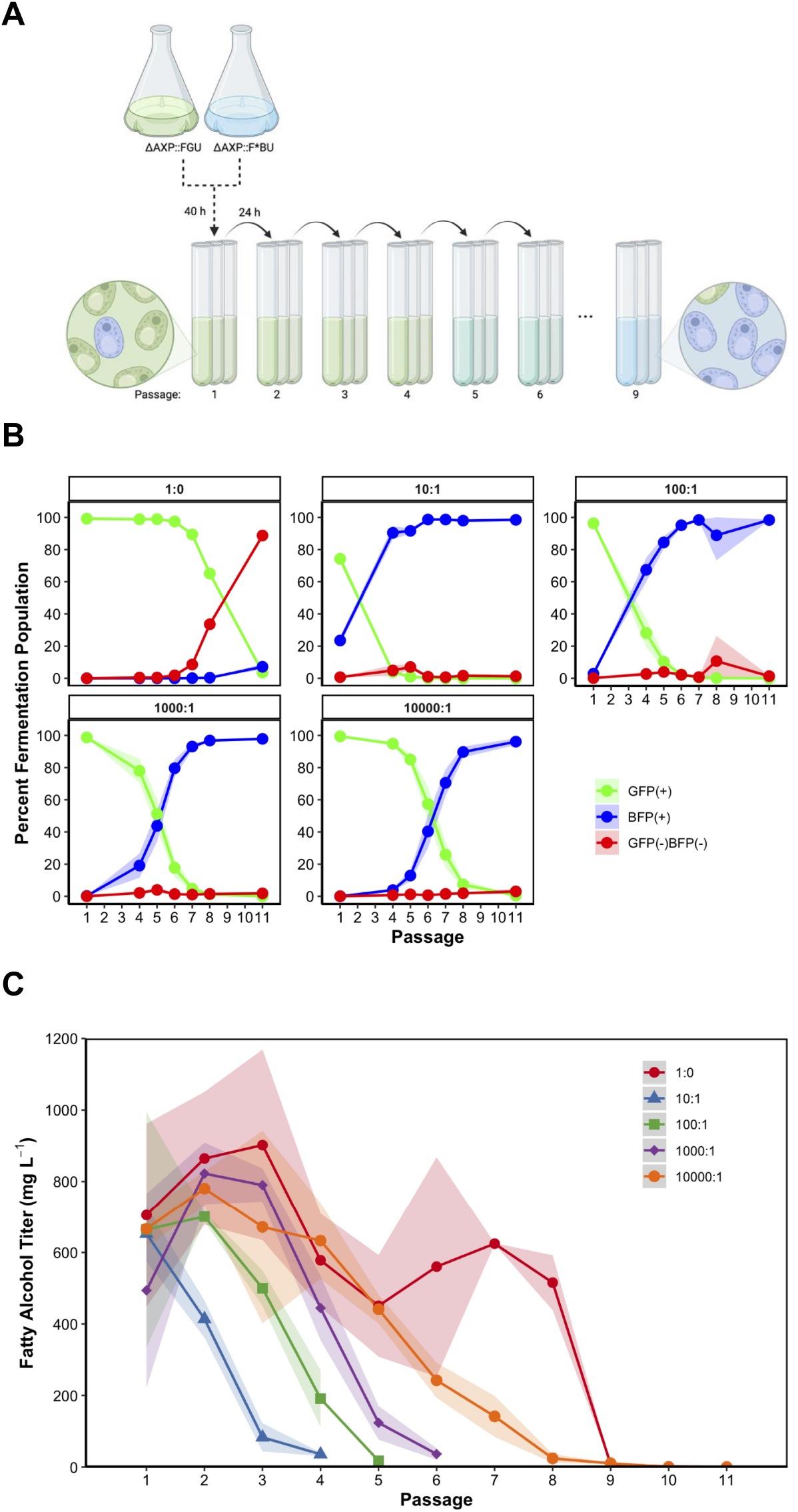
Assessing competition between fatty alcohol producers and non-producers in serial fermentation. A) Different ratios of fatty alcohol-producing (Δ*AXP::FGU*) and mutant non-producing (Δ*AXP::F*BU*) strains were co-cultured and serially passaged every 24 h. The producing strain had an initial OD600 of 0.2. B) Higher starting ratios (GFP-to-BFP ratio) of the non-producing subpopulation correlated with shortened production of fatty alcohol. C) Flow cytometric analysis to monitor the producing (green) and mutant non-producing (blue) subpopulations within the fermentation culture. Results showed the emergence of a non-producing subpopulation (red) that is non-expressing. For all sample points, the mean is shown with error bars representing standard deviation of the mean from three (n=3) biological replicates.

## 3. Discussion

Genetic instability remains a key obstacle to realizing the industrial potential of microbial fatty alcohol production. In engineered *Y. lipolytica* strains, the high metabolic burden and toxicity associated with fatty alcohol biosynthesis impose significant selective pressures that drive the emergence of low-producing or non-producing variants. Our findings demonstrate that these pressures can rapidly re-shape fermentation populations, resulting in production decline over time. Through serial passaging experiments, we observed that fatty alcohol titers quickly declined in engineered strains despite stable growth, suggesting that the production loss was not due to impaired cell growth but rather to production-linked genetic instability. It is also possible that variants with growth defects were out-competed and thus under-represented in the evolving population. The emergence of loss-of-function mutations highlights the vulnerability of engineered pathways to evolutionary dynamics, particularly when the fitness cost of production outweighs its benefits in the absence of external selection.

To address this instability, we employed a protein-fusion strategy that coupled fatty alcohol biosynthesis with essential metabolic function. Specifically, by fusing the fatty acyl-CoA reductase (FAR) to the essential glycolytic enzyme PGK1, we sought to eliminate the selective advantage of FAR inactivation. This design modestly extended production stability by one passage, while the control construct fusing FAR to GFP unexpectedly enhanced stability by four passages. Further investigation is needed to determine whether the fusion alters enzyme stability, folding, or activity, and how the native FAR compares to the two fusion proteins in terms of susceptibility to mutation or protein degradation. Understanding these differences may shed light on the mechanisms contributing to the improved production longevity observed in the fusion constructs. Despite these improvements, all strains eventually exhibited production loss, highlighting the persistent challenge of maintaining engineered function over time.

Deep sequencing of the *FAR* expression cassette across successive passages provided a detailed view of mutational events contributing to production decline. We identified a range of variants—including SNPs, frameshifts, and active site mutations—that accumulated over time and corresponded with diminished fatty alcohol titers. Notably, some mutations resulted in complete pathway inactivation, while others produced phenotypes that led to diminished, but not abolished, fatty alcohol production. Functional reconstruction of these variants revealed a spectrum of production capabilities, with certain point mutations enhancing output but likely at the cost of long-term stability. In addition, colony-level analysis demonstrated that production loss could occur even in the absence of mutations within the *FAR* cassette, suggesting extragenic or regulatory changes may also drive instability.

Our competitive co-culture experiments further demonstrated how rapidly non-producing variants can overtake productive strains in mixed populations. When non-producers were seeded at frequencies as low as10^*−*5^, they dominated the culture within six passages, and within two passages when initially seeded at 10%. These findings underscore the vulnerability of engineered production strains to evolutionary dynamics within the fermentation environment, where even small subpopulations of evolutionarily favorable variants can dramatically shift the fermentation’s productivity over time.

Collectively, this work illustrates how production loss in *Y. lipolytica* arises from a combination of genetic mutations, selective pressures, and population-level dynamics. Although strategies like gene fusion and growth-coupling can delay the onset of instability, they do not fully eliminate the evolutionary pressures that favor loss-of-function events. These findings highlight the importance of considering not only pathway optimization, but also the broader evolutionary and physiological factors that influence strain stability. By characterizing mutation patterns, production-linked selection dynamics, and population shifts over time, this study highlights key vulnerabilities in strain stability and sets the stage for future efforts aimed at mitigating stress and limiting the emergence of non-producing variants during extended cultivation. As *Y. lipolytica* continues to emerge as a platform for fatty acid-derived chemical production, addressing the underlying causes of genetic instability will be critical for enabling scalable, high-yield fermentation processes.

## 4. Conclusion

This study demonstrates that genetic instability and population heterogeneity pose significant barriers to the sustained production of fatty alcohols in engineered *Y. lipolytica*. Through serial passaging, deep sequencing, and competitive co-culture experiments, we identified how specific mutations, fitness trade-offs and subpopulation dynamics contribute to production loss over time. While fusion-based strategies—such as linking FAR to essential or fluorescent proteins—can delay the onset of instability, these interventions alone are insufficient to fully preserve strain performance across extended cultivation.

Future work should critically examine the relationship between strain productivity and genetic stability under scaled-up fermentation conditions. While high-producing strains are commercially desirable, their genetic stability may be compromised by the metabolic burden of increased production and fatty alcohol toxicity. Resolving this trade-off will require detailed investigations into how elevated production levels impact cellular metabolism and long-term genetic stability. Specifically, research should focus on developing engineering strategies that can simultaneously maintain high productivity and genetic robustness, potentially through integrating metabolic pathway optimization with mechanisms to mitigate cellular stress. A deeper understanding of the molecular mechanisms underlying metabolic burden and fatty alcohol toxicity will be crucial for developing more rational engineering approaches that can systematically address the complex challenges of industrial-scale fatty alcohol production. Understanding these fundamental molecular mechanisms will be critical for developing industrially viable strains capable of sustained, high-yield fatty alcohol production.

## 5. Materials and Methods

### 5.1. Plasmid construction

Recombinant plasmids were constructed using HiFi Assembly according to the manufacturer’s protocol, with 2 µL of assembled product used for *E. coli* transformation (New England BioLabs). NEB 10-beta Competent *E. coli* cells were used for cloning and plasmid propagation (New England BioLabs). *E. coli* plasmids were isolated using the QIAprep Spin Plasmid Mini-prep Kit (Qiagen). The genomic DNA (gDNA) of yeasts needed for the polymerase chain reaction (PCR) amplification of endogenous genes was extracted using the Wizard gDNA purification kit (Promega). Heterologous genes were codon optimized using the GenSmart™ Codon Optimization tool (GenScript). Thermo Fisher Scientific FastDigest enzymes were used for restriction digestion, and PCR amplifications were performed using Q5 high-fidelity DNA polymerase (New England BioLabs). Single-stranded DNA oligos (15-60 nucleotides) for PCR and gBlock gene fragments were synthesized by IDT. Gene fragments and clonal genes were synthesized by GenScript and Twist Bioscience, respectively. DNA was separated using gel electrophoresis, visualized using Image Lab 5.2.1 (BioRad) and subsequently purified with the Zymoclean Gel DNA Recovery Kit (Zymo Research Corporation). The LINEAR CRISPR platform was used for gene integration or knock-out (Ploessl et al., 2022). CHOPCHOP v3 was used for the specific single guide RNA (sgRNA) design (Labun et al., 2019). Whole plasmid sequencing via Oxford Nanopore sequencing was used to confirm all plasmids (Plasmidsaurus). The strains used in this study are listed in Supplementary Table S1. Plasmids and cloning primers used in this study are listed in Supplementary Table S2 and Supplementary Table S3, respectively.

### 5.2. Strain construction

*Y. lipolytica* PO1f (ATCC # MYA-2613), a leucine and uracil auxotroph devoid of extracellular proteases (Madzak et al., 2000), was used as the host strain for all studies and cultivated at 30ºC under constant agitation (250 rpm). LINEAR cassettes were obtained via restriction digest of their associated plasmids. Yeast strains were transformed with 200-500 ng of a gel-purified fragment by electroporation, as previously described (Shao et al., 2009), with a voltage of 1.8 kV mm^-1^ and plated on a selective medium corresponding to the selection marker on the LINEAR fragment and incubated at 30ºC until sizeable colonies were obtained (about 48 h). Colonies were randomly selected and plated on YPAD to isolate a pure genotype, filtering out false positives associated with extrachromosomal-based selection marker expression, and single colonies were subsequently plated on selective medium to verify the correct genotype. After 48 h of incubation, single colonies from the selective medium plate were cultured in liquid YPAD for 24 h for gDNA isolation. Loci of interest were PCR amplified and gels used to evaluate the success of the genome editing performed. Purified PCR products were sequenced using Oxford Nanopore sequencing (Plasmidsaurus) to determine the desired genome modifications).

For subsequent genome edits, the *URA3* selection marker was recovered via the site-specific CreA-*loxP* recombination system for deletion of the marker to allow recycling or other strain manipulation [123]. The plasmid p3902 CreA-Leu2 was gifted from Suzanne Sandmeyer (University of California, Irvine) and transformed into the *Y. lipolytica* strains. Cells were plated onto selective medium and incubated at 30ºC for 72 h. After incubation, a single colony was randomly selected and inoculated in liquid SC-LEU to recycle *URA3*. After 72 h of cultivation, the saturated culture was diluted 10-100 times, and the diluted cells were plated on SC-LEU+5-FOA and incubated until sizeable colonies were obtained (about five to seven days). After incubation, a single colony was randomly selected and simultaneously inoculated in liquid YPAD and SC-URA to lose the Cre plasmid and verify the removal of *URA3*. After culturing for 48 h, the saturated YPAD culture was diluted 100-1000 times, and the diluted cells were plated on YPAD. After 72 h of incubation, a single colony was simultaneously inoculated in liquid YPAD and SC-URA to ensure the Cre plasmid has been lost and *URA3* successfully removed.

### 5.3. Media and culture conditions

Selective medium used to grow transformants contained complete supplement mixture (CSM) with appropriate amino acid dropouts (added according to the manufacturing instructions), 1.67 g/L yeast nitrogen base without amino acids and ammonium sulfate, 5 g/L ammonium sulfate, 43 mg/L adenine hemisulfate and 20 g/L dextrose. Nonselective medium contained 10 g/L yeast extract, 20 g/L peptone, 100 mg/L adenine hemisulfate and 20 g/L dextrose. For fatty alcohol production cultures, the liquid medium contained 40 g/L dextrose. All strains were cultured at 30ºC with shaking at 250 rpm.

### 5.4. Growth characterization of fatty alcohol-producing strains

Seed cultures of *Y. lipolytica* strains were grown in 25-mL selective medium (containing 40 g/L dextrose and appropriate amino acids dropout) within a 250-mL baffled flask for 40 h and subsequently inoculated into 25-mL nonselective medium (containing 40 g/L dextrose) within a 250-mL baffled flask to a starting OD_600_ of about 0.2. Cells were cultured for 120 h. For the first 12 h, the OD_600_ was measured every 3 h. Afterwards, the OD_600_ was measured every 12 h.

### 5.5. Serial passaging and fermentation of fatty alcohol-producing strains

To assess fatty alcohol production stability, fatty alcohol-producing *Y. lipolytica* strains were pre-cultured, plating on selective medium and inoculating in 25-mL selective medium (containing 40 g/L dextrose and appropriate amino acid dropouts) within a 250-mL baffled flask for 40 h. The seed culture was subsequently inoculated into 6-mL nonselective medium within a 55-mL round bottom borosilicate glass tube to achieve an initial OD_600_ of about 0.2 and cultured for 120 h (Passage 1). After 24 h, 1% of the fermentation culture was transferred, or passaged, into 6-mL of fresh medium and cultured for 120 h (Passage 2). Passaging continued with each subsequent culture for a total of nine passages.

For assessing the population dynamics of cultures containing non-producers, the seeding ratio (producing-to-non-producing strain with an initial OD600 of about 0.2 for the producer) of the starting culture (Passage 1) varied between 1:0 (no nonproducer added) to 10,000:1. For this study, the strains Δ*AXP::FGU* (producing strain) and Δ*AXP::F*BU* (non-producing strain) were used. Serial passaging and fermentation proceeded as previously described.

### 5.6. Quantification of fatty alcohol production via GC-MS analysis

Fatty alcohols were extracted from a 0.5 mL aliquot of a 6-mL fermentation culture by addition of 0.5 mL ethyl acetate, spiked with 50 µg tridecanol (C13-ol) as an internal standard. Glass beads (acid-washed, 425-600 µm) were added to the microcentrifuge tube to promote disruption of cells during extraction. The mixture was vortexed at 2,500 rpm for 20 minutes and subsequently centrifuged at 14,800 rpm for 10 minutes to separate the organic layer. An aliquot of 100 µL from the organic layer was dried (either under nitrogen or air dried overnight) and silylated with N,O-Bis(trimethylsilyl)trifluoroacetamide (BSFTA) reagent by resuspending the dried sample in 100 µL of BSFTA and incubating at 60ºC for 30 minutes. Samples for GC-MS analysis of fatty alcohol content were analyzed on an Agilent Technologies Model 7890A GC coupled to a 5975C Mass Spectrometry detector capable of electrical ionization.

### 5.7. Flow cytometric analysis for quantification of GFP and BFP expression

After 48 h, an aliquot was taken from the fermentation culture, washed and diluted 20-fold with 1X PBS (pH 7.4) Samples were analyzed for GFP and/or BFP fluorescence intensity at 488 nm and 399 nm, respectively, using a FACSCanto flow cytometer (BD Biosciences). The fluorescence intensity distribution of GFP and BFP-expressing cells was calculated using BD FACSDiva 8.0.1. Data for forward scatter (FSC) and side scatter (SSC) were collected in logarithmic mode using instrument voltage settings typical for yeast cell populations. Fluorescence intensities were quantitatively measured in logarithmic mode, and values were recorded as mean fluorescence intensity for each event. Initial gating to identify yeast cells was based on FSC and SSC data analyzed in dual-parameter dot plots, using data from control samples containing suspended non-expressing yeast cells to distinguish between background/noise and actual yeast cell events. Most background/noise events were eliminated by setting an SSC acquisition threshold. Gated yeast cell fluorescence data were analyzed in dot plots of SSC versus the relevant fluorescent parameter, either GFP or BFP.

### 5.8. Sequencing, quality control, and read pre-processing

Yeast populations harvested after serial passaging were sequenced using an Illumina MiSeq platform, generating 250 bp paired-end reads across two sequencing lanes. Initial quality assessment was performed with FastQC v0.11.9, with lane-specific reports aggregated via MultiQC v1.19 to evaluate base quality scores, adapter contamination, and read length distributions. Adapter sequences were trimmed using BBDuk (BBMap v39.01) with a 23-mer seed length (minimum k = 11, Hamming distance = 1, with pair-overlap detection enabled). Quality filtering was applied at both 5’ and 3’ ends using a stringent Q30 threshold, and reads shorter than 25 bp post-trimming were discarded. The processed reads from both lanes were subsequently merged by sample, with quality improvements verified through a second round of FastQC/MultiQC analysis.

### 5.9. Reference preparation, mapping, variant calling, and mutation analysis

Three construct-specific reference sequences corresponding to FAR, FARGFP, and FARPGK1 experimental groups were indexed using samtools faidx v1.20 and processed through the GATK-flow pipeline (isugifNF, 2024), a custom Nextflow-based implementation of GATK best practices. The pipeline orchestrated read alignment, post-alignment processing, and variant discovery using GATK HaplotypeCaller v4.4.0.0, generating VCF files containing single-nucleotide variants (SNVs) and insertions/deletions (INDELs). The resulting alignment files were converted from CRAM to sorted, indexed BAM format using samtools v1.20 to facilitate downstream analyses. Nucleotide frequency calculations at each genomic position were performed using bam-readcount v1.0.1 (Khanna et al., 2022). Subsequent analytical steps quantified SNV and INDEL frequencies across the genome, characterized mutation distribution by genomic feature, and monitored allele frequency trajectories across sequential passages to identify genomic loci potentially subject to positive selection.

## CRediT authorship contribution statement

**Jeniffer Perea-Lopez**: Writing – review & editing, Writing – original draft, Conceptualization, Methodology, Investigation, Formal Analysis, Visualization. **Yuxin Zhao**: Methodology, Investigation, Conceptualization. **Viswanathan Satheesh**: Methodology, Software, Data Curation. **Zhanyi Yao**: Investigation. **Dan Chen**: Investigation. **Zengyi Shao**: Writing – review & editing, Supervision, Conceptualization, Project administration, Funding acquisition.

## Acknowledgments

All figures were created with BioRender. This work was funded by the DOE Center for Advanced Bioenergy and Bioproducts Innovation (U.S. Department of Energy, Office of Science, Biological and Environmental Research Program under Award Number DE-SC0018420). Any opinions, findings, and conclusions or recommendations expressed in this publication are those of the author(s) and do not necessarily reflect the views of the U.S. Department of Energy.

