## Supplementary Information for "Developing engineering strategies to enhance the genetic stability of fatty alcohol-producing strains for production scale-up"

Supplementary Table S1: Strains used in this study.

| Strain name | Genotype | Description | Reference |
|---|---|---|---|
| *Y. lipolytica* PO1f | *MATa, leu2-270, ura3-302, xpr2-322, axp-2* | | |
| ΔAXP::FAR | *axp::FAR-GFP-loxP-URA3-loxP* | Integration of *FAR* expression cassette at the AXP locus | Zhao et al. |
| ΔAXP::FPGU | *axp::FAR-(GGGGS)$_3$-PGK1-GFP-loxP-URA3-loxP* | *FAR-PGK1* fusion cassette integrated at the AXP locus; *PGK1* includes modified N20 sequence | This study |
| ΔAXP::FPG | *axp::FAR-(GGGGS)$_3$-PGK1-GFP; pgk1::loxP-URA3-loxP* | Derived from ΔAXP::FPGU; deletion of native *PGK1* | This study |
| ΔAXP::FGU | *axp::FAR-(GGGGS)$_3$-GFP-loxP-URA3-loxP* | Control strain expressing *FAR*-GFP fusion cassette | This study |
| ΔAXP::F*BU | *axp::FAR(Y248F)-(GGGGS)$_3$-mTagBFP2-loxP-URA3-loxP* | Non-producing control strain with inactive *FAR* mutant (Y248F) fused to BFP | This study |

Supplementary Table S2: Plasmids used in this study.

| Plasmid name | Description | Reference |
|---|---|---|
| YL-CashGem-BAT1-URA3-LINEAR | LINEAR CRISPR expression vector; used for HR-based genome editing in *Y. lipolytica* | Ploessl et al., 2022 |
| p3902 CreA-Leu2 | Episomal plasmid expressing Cre recombinase | Suzanne Sandmeyer's Lab |
| LINEAR ΔAXP::FAR-GFP-URA3 | For integration of *FAR* expression cassette at the AXP locus with *URA3* selection and GFP reporter | Zhao et al. |
| LINEAR ΔAXP::MTSroGFP-LEU2 | For mitochondrial-targeted roGFP (MTSroGFP) integration at AXP locus with *LEU2* selection | Yao et al. |
| LINEAR ΔB1::FAR-$(GGGGS)_3$-GFP-URA3 | For integration of *FAR*-GFP fusion cassette at the B1 locus with *URA3* selection and GFP reporter | Yao et al. |
| LINEAR ΔPGK1::FAR-$(GGGGS)_3$-PGK1-URA3 | For integration of *FAR-PGK1* fusion cassette at the *PGK1* locus with *URA3* selection; *PGK1* includes modified N20 sequence | This study |
| LINEAR ΔAXP::FAR-$(GGGGS)_3$-PGK1-GFP-URA3 | For integration of *FAR-PGK1* fusion cassette at AXP locus with *URA3* selection and GFP reporter | This study |
| LINEAR ΔPGK1::HPT | For knockout of native *PGK1* with hygromycin resistance selection | This study |
| LINEAR ΔPGK1::HPT v2 | For knockout of native *PGK1* with hygromycin resistance selection; targeting of *PGK1* intronic region | This study |
| LINEAR ΔPGK1::URA3 | For knockout of native *PGK1* with *URA3* selection; targeting of *PGK1* intronic region | This study |
| LINEAR ΔAXP::FAR-$(GGGGS)_3$-GFP-URA3 | For integration of *FAR*-GFP fusion cassette at AXP locus with *URA3* selection | This study |
| LINEAR ΔAXP::FAR(Y248F)-$(GGGGS)_3$-BFP-URA3 | For integration of catalytically inactive *FAR* (Y248F) fused to BFP at AXP locus with *URA3* selection | This study |
| LINEAR ΔB1::FAR-$(GGGGS)_3$-GFP-LEU2 | For B1 locus integration of *FAR*-GFP fusion cassette with *LEU2* selection | This study |

Supplementary Table S3: Primers and DNA fragments used in this study.

| Primer | Sequence (5′-to-3′) | Use |
|---|---|---|
| *Plasmid LINEAR ΔPGK1::FAR-(GGGGS)$_3$-PGK1-URA3* | | |
| S1 | GCTCATTCTCCCCGGCGCTCTCCGATCTC TTCCGACGAAAATGGCCATCCAGCAGGTC | Amplify pTEF1in-FAR |
| S2 | TTTGAGATGCGTGTGAAATGTGCTCATCG ACCCTCCGCCTCCGCTACCGCCTCCACCA GAGCCTCCTCCACCGGCGGCCTTCTTCCG CTG | Amplify pTEF1in-FAR (with (GGGGS)$_3$ tail) |
| S3 | GGTGAGGTCATTCTCCTCGAGAACCTCCG ATTTCACCCTGAGGAGGAGGGATCCCACA AG | Amplify second half of PGK1 |
| S4 | GCACACAGAACCGGGCACTCACTTCCCCA TCCACACTTCCTTACTTCTTCTCGGAGAG AG | Amplify second half of PGK1 |
| S5 | AGACCCTTCCCGGTGTCGCTGCTCTCTCC GAGAAGAAGTAAGGAAGTGTGGATGGGG AAG | Amplify pex20t |
| S6 | AGCATACATTATACGAAGTTATAGGTAAC GGCTTGGGTGCTATATTTGACGATTGACG C | Amplify pex20t |
| S7 | TATTCATTCATGTTAGTTGCGTCAATCGT CAAATATAGCACCCAAGCCGTTACCTATA AC | Amplify *URA3* cassette |
| S8 | ATCAACGTTATATTCCTCATTACACATGC TGTAAATAGCCTCGCTTCTCTCGATAACT TC | Amplify *URA3* cassette |
| S9 | CGCGCGTAATACGACTCACTATAGGGCGA ATTGGGTACCGGTTTGGCATTGAATATTC AG | Amplify homology arms upstream of PGK1 |
| S10 | GGAGGAGGAGGTGTCGGCGTGGTGGACC TGCTGGATGGCCATTTTCGTCGGAAGAG ATCG | Amplify homology arms upstream of PGK1 |
| S11 | TATAATGTATGCTATACGAAGTTATCGAG AGAAGCGAGGCTATTTACAGCATGTGTAA TG | Amplify homology arms downstream of PGK1 |
| S12 | CTCAGAGCCTCGGCCCAGCCTTCGGCCCT TTTGGGTTTGACTACTCCTGTGTAACG | Amplify homology arms downstream of PGK1 |

| Primer | Sequence (5′-to-3′) | Use |
|---|---|---|
| S14 | CACTGGCCGGTCGATAATTTAAC | Amplify mig1t |
| PGK1 | TGAAGACCGCCCGGCAGCGGAAGAAGGC CGCCGGTGGAGGAGGCTCTGGTGGAGGC GGTAGCGGAGGCGGAGGGTCGATGAGCA CATTTCACACGCATCTCAAAATGTCTCTT ACCAACAAGCTCTCCATCAAGGATCTCGA TCTCAAGAACAAGCGAGTCTTCATCCGAG TCGACTTTAACGTTCCTCTCGATGGCACC ACCATCACCAACAACCAGCGAATTGTTGC TGCCCTGCCCTCCATCAAGTATGCCATTG ATCAGGGTGCCAAGGCTGTGATCCTTGCT TCTCATCTCGGCCGGCCCAACGGCCAGCG AGTCGAGAAGTACTCTCTCAAGCCCGTCC AGGCCGAGCTTTCAAAGCTCCTTGGCAAG CCCGTCACCTTCCTTGACGACTGCGTCGG CCCCAAGGTTGAGGAGGAGGTCTCCAAG GCCAAGGACGGTGAGGTCATTCTCCTCGA GAACCTCCGATTTCACCCTGAGGAGGAGG GATCCCACAAG | DNA fragment containing $(GGGGS)_3$ and first half of PGK1 |
| PGK1_N20 | GGTAGGAAAATATTATGTCTATACGAGAC GAATACGGTCACCGACTCTAATGTAAGTG ATACCATGTATAGTAGTGTCTTCTGGTAC AGAACTCCAAGTAATGTGTACTTAATCCC CTGTAATACAATACCGTGCTCTCATTCTC TTACGGTATCATTACATTCATCTATATAC AGCCAATGGGCAATGCGTGTGCTGAGCT CAGCACGTTAAATTATCGACCGGCCAGTG GTCCCATTCGCCATGCCGAAGCATGTTGC CCAGCCGGCGCCAGCGAGGAGGCTGGGA CCATGCCGGCCAAAAGCACCGACTCGGTG CCACTTTTTCAAGTTGATAACGGACTAGC CTTATTTTAACTTGCTATTTCTAGCTCTA AAACCGGGGTGGAATCGCAGGTTCGACG AGCTTACTCGTTTCGTCCTCACGGACTCA TCAGGAACCTCTGCGGTTAGTACTGCAAA AAGTGCTGGTCGGATGACGTGGCGTCTT GTGTCGATGGG | DNA fragment containing sgRNA cassette for PGK1 knockout |
| *Plasmid LINEAR ΔAXP::FAR-$(GGGGS)_3$-PGK1-GFP-URA3* | | |
| S15 | CGTCCTTGTCCAAAGTTTGAGGTACCAGA GACCGGGTTGGCGGC | Amplify pTEF1in-FAR-$(GGGGS)_3$-PGK1-pex20t |



*Continued on next page*

| Primer | Sequence (5′-to-3′) | Use |
|---|---|---|
| S16 | AAAAAACGGGCGCCAAACTCCGCCGGCG GCTATATTTGACGATTGACGCAAC | Amplify pTEF1in-FAR-(GGGGS)$_3$-PGK1-pex20t |
| *Plasmid LINEAR ΔPGK1::HPT* | | |
| S17 | CCTCATTACACATGCTGTAAATAGCGGTA CCATCTAGAACGCCGCCTATG | Amplify HPT cassette |
| S18 | CGCTCTCCGATCTCTTCCGACGAAAGGTA CCGCATTAGTACGATTCGAGC | Amplify HPT cassette |
| S19 | TTATATGCTCGAATCGTACTAATGCGGTA CCTTTCGTCGGAAGAGATCG | Amplify homology arms upstream of PGK1 |
| S20 | TACGACTCACTATAGGGCGAATTGGGGTT TGGCATTGAATATTCAG | Amplify homology arms upstream of PGK1 |
| S21 | AATCTACGCTTGTTCAGACTTTGTACTAG TTTCTTTGTCTGGC | Amplify sgRNA cassette for PGK1 knockout and homology arm downstream of PGK1 |
| S22 | AGTTATCATAGGCGGCGTTCTAGATGGTA CCGCTATTTACAGCATGTGTAATG | Amplify sgRNA cassette for PGK1 knockout and homology arm downstream of PGK1 |
| *Plasmid LINEAR ΔPGK1::HPT v2* | | |
| S23 | AATTTAACGTGCTGAGCTC | Amplify mig1t and homology arm downstream of PGK1; pair with primer S22 |

*Continued on next page*

| Primer | Sequence (5′-to-3′) | Use |
|---|---|---|
| PGK1_intron_N20 | CTTTAGCCAAGGGTATAAAAGACCACCGTCCCCGAATTACCTTTCCTCTTCTTTTCTCTCTCTCCTTGTCAACTCACACCCGAAATCGTTAAGCATTTCCTTCTGAGTATAAGAATCATTCAAAATGGTGAGTTTCAGAGGCAGCAGCAATTGCCACGGGCTTTGAGCACACGGCCGGGTGTGGTCCCATTCCCATCGACACAAGACGCCACGTCATCCGACCAGCACTTTTTGCAGTACTAACCGCAGTTGTGTCTGATGAGTCCGTGAGGACGAAACGAGTAAGCTCGTCACACAACTGATTTCGAAACGGTTTTAGAGCTAGAAATAGCAAGTTAAAATAAGGCTAGTCCGTTATCAACTTGAAAAAGTGGCACCGAGTCGGTGCTTTTGGCCGGCATGGTCCCAGCCTCCTCGCTGGCGCCGGCTGGGCAACATGCTTCGGCATGGCGAATGGGACCACTGGCCGGTCGATAATTTAACGTGCTGAGCTCAGCACA | DNA fragment containing sgRNA cassette for PGK1 knockout targeting the intronic region |
| *Plasmid LINEAR ΔPGK1::URA3* | | |
| S24 | CCTCATTACACATGCTGTAAATAGCGGTACCCTCGCTTCTCTCGATAACTTC | Amplify *URA3* cassette |
| S25 | CGCTCTCCGATCTCTTCCGACGAAAGGTACCACCCAAGCCGTTACCTATAAC | Amplify *URA3* cassette |
| *Plasmid LINEAR ΔAXP::FAR-(GGGGS)$_3$-GFP-URA3* | | |
| S26 | GTAAAGAGTGATAAATAGCGTTAACCGCCGGCGGCTATATTTGACGATTGACGCAAC | Amplify pTEF1in-FAR-(GGGGS)$_3$-PGK1-pex20t; pair with primer S15 |
| *Plasmid LINEAR ΔAXP::FAR-(GGGGS)$_3$-GFP-URA3* | | |
| S27 | CGACCCTCCGCCTCCGCTACCGCCTCCACCAGAGCCTCCTCCACCGGCGGCCTTCTTCCGCTG | Amplify pTEF1in-FAR- GGGGS)$_3$; pair with primer S15 |

*Continued on next page*

| Primer | Sequence (5′-to-3′) | Use |
|---|---|---|
| S28 | GTAGCGGAGGCGGAGGGTCGATGGTGAG CAAGCAGATCCTGAAG | Amplify GFP |
| S29 | CACTCACTTCCCCATCCACACTTCCTTAC ACCCACTCGTGCAGGCT | Amplify GFP |
| S30 | GGAAGTGTGGATGGGGAAGTG | Amplify pex20t |
| S31 | TGTAAAGAGTGATAAATAGCGTTAACCGC CGGCGGCTATATTTGACGATTGACGCAAC | Amplify pex20t |
| *Plasmid LINEAR ΔAXP::FAR(Y248F)-(GGGGS)$_3$-BFP-URA3* | | |
| S32 | CCCAGCCACTTGGTGAAGGTGAAGGTGT CGGACCAGCCATAC | Amplify pTEF1in and first half of FAR(Y248F); pair with primer S15 |
| S33 | GGCATCCGGGAGGCCAACCGGTATGGCT GGTCCGACACCTTC | Amplify second half of FAR(Y48F) and (GGGGS)$_3$ |
| S34 | CGACCCTCCGCCTCCGCTACCGCCTCCAC CAGAGCCTCCTCCACCGGCGGCCTTCTTC CGCTG | Amplify second half of FAR(Y48F) and (GGGGS)$_3$ |
| S35 | GTAGCGGAGGCGGAGGGTCGATGTCTGA GCTGATCAAGGAG | Amplify mTagBFP2 |
| S36 | ACTTCCCCATCCACACTTCCTCAATTGAG CTTGTGGCCCAG | Amplify mTagBFP2 |

*Continued on next page*

| Primer | Sequence (5′-to-3′) | Use |
|---|---|---|
| *Yl*mTagBFP2 | ATGTCTGAGCTGATCAAGGAGAACATGC<br>ACATGAAGTTGTACATGGAGGGTACCGT<br>TGACAACCATCATTTTAAGTGCACCAGCG<br>AAGGCGAGGGAAAGCCCTACGAGGGCAC<br>GCAAACCATGCGAATCAAAGTGGTCGAA<br>GGAGGACCGCTCCCCTTTGCCTTTGACAT<br>TCTTGCCACTTCTTTTCTCTACGGCTCCA<br>AGACTTTCATCAACCACACCCAGGGTATT<br>CCCGACTTCTTCAAGCAGTCGTTCCCCGA<br>GGGCTTCACCTGGGAGCGAGTGACAACG<br>TATGAAGACGGAGGAGTGCTAACGGCGA<br>CACAAGATACCTCGCTGCAGGACGGGTGT<br>CTCATCTACAACGTCAAGATTCGGGGTGT<br>CAACTTCACTTCCAACGGACCTGTCATGC<br>AGAAGAAAACTTTGGGCTGGGAAGCCTT<br>CACAGAGACCCTGTACCCCGCAGATGGGG<br>GTCTTGAGGGACGTAACGACATGGCTCTC<br>AAGCTTGTTGGTGGCTCACATCTGATTGC<br>AAATGCCAAGACCACCTATAGATCTAAGA<br>AGCCTGCTAAGAACCTGAAGATGCCTGTC<br>TACTACGTGGACTACCGACTGGAACGCAA<br>AGAGGCCAACAATGAGACTTATGTAGAG<br>CAGCACGAGGTGGCTGTTGCTCGATACTG<br>CGATCTGCCAAGTAAACTGGGCCACAAGC<br>TCAATTGA | Codon optimized mTagBFP2 |
| *Plasmid LINEAR ΔB1::FAR-(GGGGS)$_3$-GFP-LEU2* | | |
| S37 | CGTCAATCGTCAAATATAGCCGCCGGCGT<br>GAAGCGCTCGTGATTGT | Amplify *LEU2* cassette |
| S38 | GTAGAAAATCGCCAAGTGGATTAATTAAC<br>ACTAACCTACCAAACACCAC | Amplify *LEU2* cassette |
